# Oxidative stress and vacuolar variants influence paraquat response in *Saccharomyces cerevisiae* of different genetic variants

**DOI:** 10.1101/2025.05.27.656361

**Authors:** Juan Carlos Rubilar, Benjamín Szenfeld, Francisco A. Cubillos, Andrés. D. Klein

## Abstract

The biological effects of paraquat (PQ), an herbicide linked to Parkinson’s disease (PD) risk in humans can be studied in *Saccharomyces cerevisiae* due to its evolutionary conservation with mammals. To understand how genetic background influences PQ toxicity, we treated four yeast strains (NA, SA, WA, and WE) with PQ and assessed physiological (growth curves), molecular (superoxide and peroxide levels) and cellular consequences (vacuolar disaggregation) on each genetic background. PQ significantly reduced specific growth rates (μMax) in WE and WA, while SA and NA remained unaffected. Furthermore, PQ increased superoxide and peroxide levels across the strains, but to different extents, being the SA and WE the most affected. PQ also influenced vacuolar morphologies strain-dependently, shifting from one large organelle to small disaggregate vacuoles, with WE being the most susceptible. Interestingly, we found an inverse association between superoxide levels and μMax. Given the known involvement of lysosomal dysfunction in pesticide-induced PD, we investigated correlations between predicted missense variants in vacuolar genes and PQ responses across the yeast strains. We identified associations between *fen2* variants, the human orthologue of *SLC17A5*, and vacuolar disaggregation. To validate this, we exposed a *fen2*-deleted strain to PQ, which exhibited increased vacuolar disaggregation. In conclusion, our findings demonstrate that the strain-specific susceptibility to PQ is associated with superoxide levels and that *fen2* likely plays a role in the vacuolar adaptive response to PQ exposure. We speculate that the most resistant strains may facilitate the development of novel therapeutics for humans exposed to PQ.

## Introduction

Paraquat (PQ) is a potent herbicide widely employed globally due to its efficacy in eradicating harmful organisms detrimental to crops (Tudi et al., 2021). Despite its widespread use, with an estimated global consumption of 3.5 million tons annually (Pretty and Bharucha, 2015), PQ poisoning causes ∼20,000 deaths yearly (Ball et al., 2019; Sharma et al., 2012). A clear link between PQ exposure and Parkinson’s disease (PD) has been established (Paul et al., 2024), raising environmental and human health concerns (Coria and Elgueta, 2022).

PD pathogenesis is complex, with the etiology of most cases remaining unknown (Ashraf et al., 2024; Morris et al., 2024). However, it is well-established that a complex interplay between genetic and environmental factors contributes to its development. Several PD genetic risk factors interact with pesticide and herbicide exposure (Fleming, 2017; Prasad et al., 2007). Nevertheless, the precise molecular mechanisms of gene-environment interaction remain elusive. Elucidating them is essential for achieving early diagnosis, preventative strategies, and translating potential therapeutic interventions (Burbulla and Krüger, 2011; Dardiotis et al., 2013).

PQ inhibits mitochondrial complex I by reducing NADH-cytochrome b5 and NADH-ubiquinone oxidoreductases, increasing superoxide anion, which is further converted into hydrogen peroxide and hydroxyl radicals (Elkholy et al., 2023; Hernandez-Baixauli et al., 2024; Torres-Rojas et al., 2020). Additionally, PQ induces the opening of the mitochondrial permeability transition pore, reducing intra-mitochondrial calcium retention capacity and decreasing the membrane potential, which also contributes to increasing reactive oxygen species (ROS) levels (See et al., 2022; Song et al., 2019). Mitochondrial damage directly impacts the lysosomal pathway via membrane contact sites, leading to decreased lysosomal enzyme activities, impaired sphingolipids degradation and autophagy blockage (Li et al., 2020; Rubilar et al., 2024). These disruptions contribute to α-synuclein (α-syn) aggregation, dopaminergic neuron death (Klein and Mazzulli, 2018; Zhang et al., 2016), and ultimately motor impairments (Islam et al., 2021).

Yeast models represents a powerful biological tool for dissecting the underlying genetic and molecular mechanisms of PD pathogenesis due to its high degree of conservation with humans (Liu et al., 2017), rapid growth rate and generation time, availability of panels of strains bearing genotypic variability, and capacity for rapid characterization and implementation of large-scale studies using yeast strains with single gene deletions (Mülleder et al., 2012; Winzeler et al., 1999).

In this study, we investigated the biological effects of PQ exposure on four *S. cerevisiae* strains with distinct genetic backgrounds. We employed a multi-tiered approach, examining its consequences at physiological (growth rates), molecular (ROS levels), and cellular (vacuolar morphology) levels and the interplay between them.

## Results

### Paraquat-induced strain-specific growth rates responses

We chose four phylogenetically distinct strains exhibiting extensive genomic variability, whose genomes are sequenced: Y12 (Sake, SA), YPS128 (North American, NA), DBVPG6044 (West African, WA) and DBVPG6765 (Wine/European, WE). As a proxy of cell health, we measured their growth rates under control and PQ exposure. Initially, we set the lowest PQ concentration, which induced reproducible variation across strains. We tested a range of 12.5 - 125 μg/mL (data not shown) and chose 75 μg/mL to discriminate between strains. Under control conditions (YNB media 2% glucose) all strains exhibited similar growth patterns (Figure 1A). However, upon PQ exposure we identified a lower growth rate in the WE background than untreated cells (susceptibility phenotype) (Figure 1B). This effect was quantified by calculating the Area Under the Curve (AUC) (p-value = 0.0013, ANOVA) (Figure 1C). Conversely, SA, NA and WA exhibit the highest tolerance to PQ (resistance phenotype, p-value = ns, ANOVA) (Figure 1B and C).

**Figure 1:**
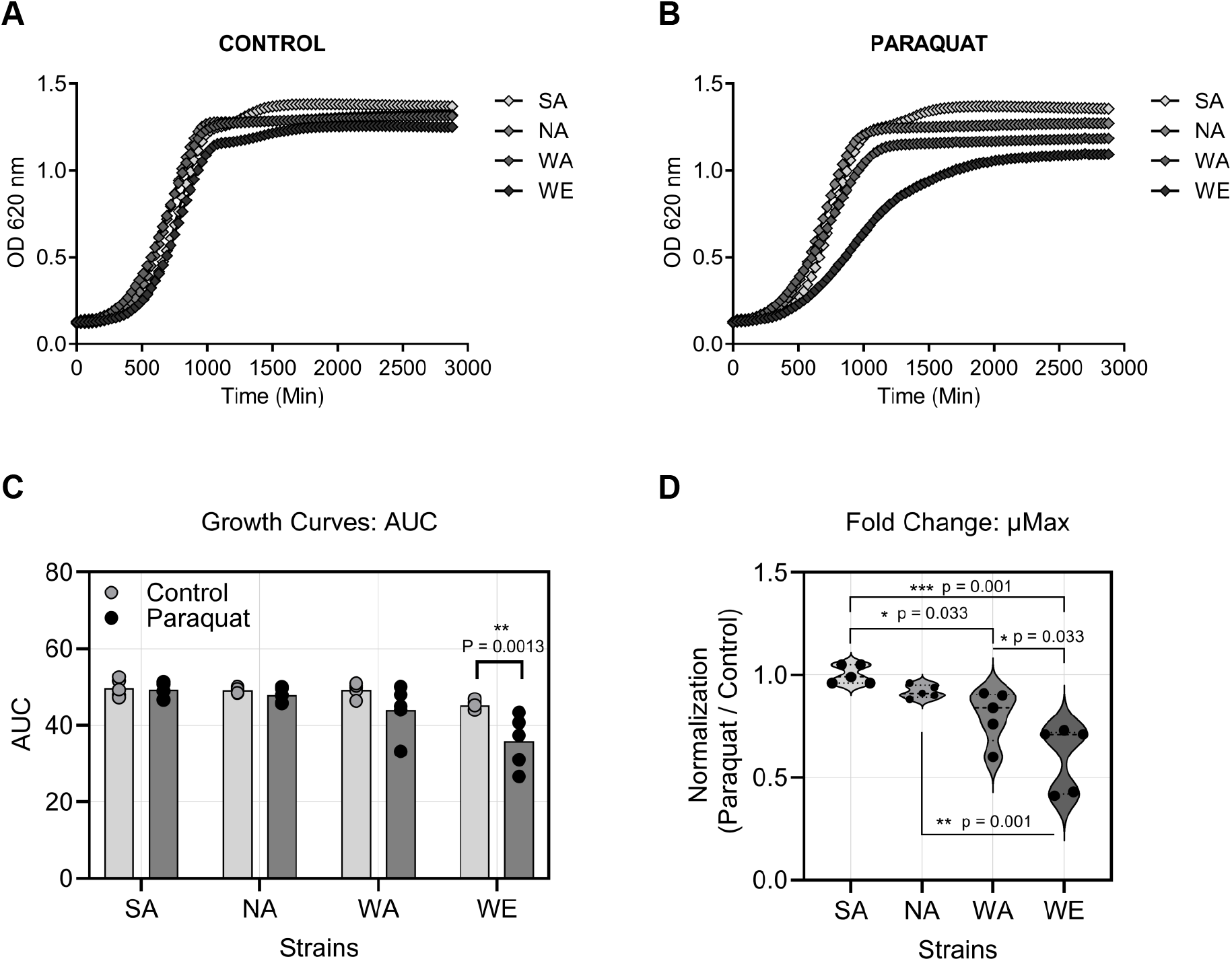
Reproductive fitness in *S. cerevisiae* parental strains against PQ exposure. (A) Growth curves of SA, NA, WA, and WE yeast strains in control condition, and (B) exposed to PQ (75 μg/mL). (C) Area under the curve (AUC). (D) Fold change of μMax (PQ/mock). Biological replicates “N= 5”, Values are mean ± SE, Shapiro Wilk normality test p-value > 0.05, ANOVA test p-value < 0.05, post-hoc Holm-Sidak (AUC and fold change). p-value ≤ 0.05 (*), p-value < 0.01 (**), p-value < 0.001 (***), p-value < 0.0001 (****), ns = not significant (p-value > 0.05). Statistical analysis was performed in SPSS software version 20.0 and plotted in GraphPad.

From the growth curves, we calculated μMax, which corresponds to the maximum rate at which the yeast can increase its population size as another proxy of reproductive fitness. PQ exposure differentially impacted μMax across the strains. PQ-treated WE exhibited a decrease in μMax compared to NA (p-value = 0.0012, ANOVA) (Figure 1D). Notably, the WE strain exhibited a 40% reduction in μMax compared to SA (p-value = 0.0001, ANOVA), with WA showing a 20% decrease when compared to SA (p-value = 0.033, ANOVA) (Figure 1D). Furthermore, we found a significant difference between WA and WE (p-value = 0.033, ANOVA). These results demonstrate that PQ differentially impacts each strain, with WE and WA exhibiting the most significant susceptibility. Therefore, PQ exposure has a strain-specific impact on yeast reproductive fitness and physiological adaptation.

### PQ induced strain-specific ROS responses

Given that PQ inhibits mitochondrial complex I, we analyzed superoxide anion levels with DHE, under control and PQ treatments (75 μg/mL for 48-hours) by microscopy. Under control conditions, we found no visible oxidation of the probe (Figure 2A). Upon PQ exposure, oxidized DHE binds to DNA, emitting visible red fluorescence (Figure 2A). All PQ-treated strains exhibited probe oxidation (red fluorescence). Interestingly, the NA strain exhibited lower fluorescence intensity compared to WA (p-value = 0.0265, ANOVA) and WE (p-value = 0.0027, ANOVA) (Figure 2B). Similarly, the SA strain showed lower intensity fluorescence compared to WA (p-value = 0.0092, ANOVA) and WE (p-value = 0.0011, ANOVA) (Figure 2B), indicating lower oxidative stress levels in these strains when were exposed to PQ (Figure 2B). In addition, PQ exposure induced significant changes in hydrogen peroxide levels when compared to control conditions (Figure 2C). Furthermore, significant differences in DCF fluorescence rates were observed among strains SA and WA after PQ treatment (p-value = 0.0279, ANOVA) (Figure 2D). These results indicate that PQ differentially impacts the yeast strains superoxide anion and hydrogen peroxide levels in the yeast strains.

**Figure 2:**
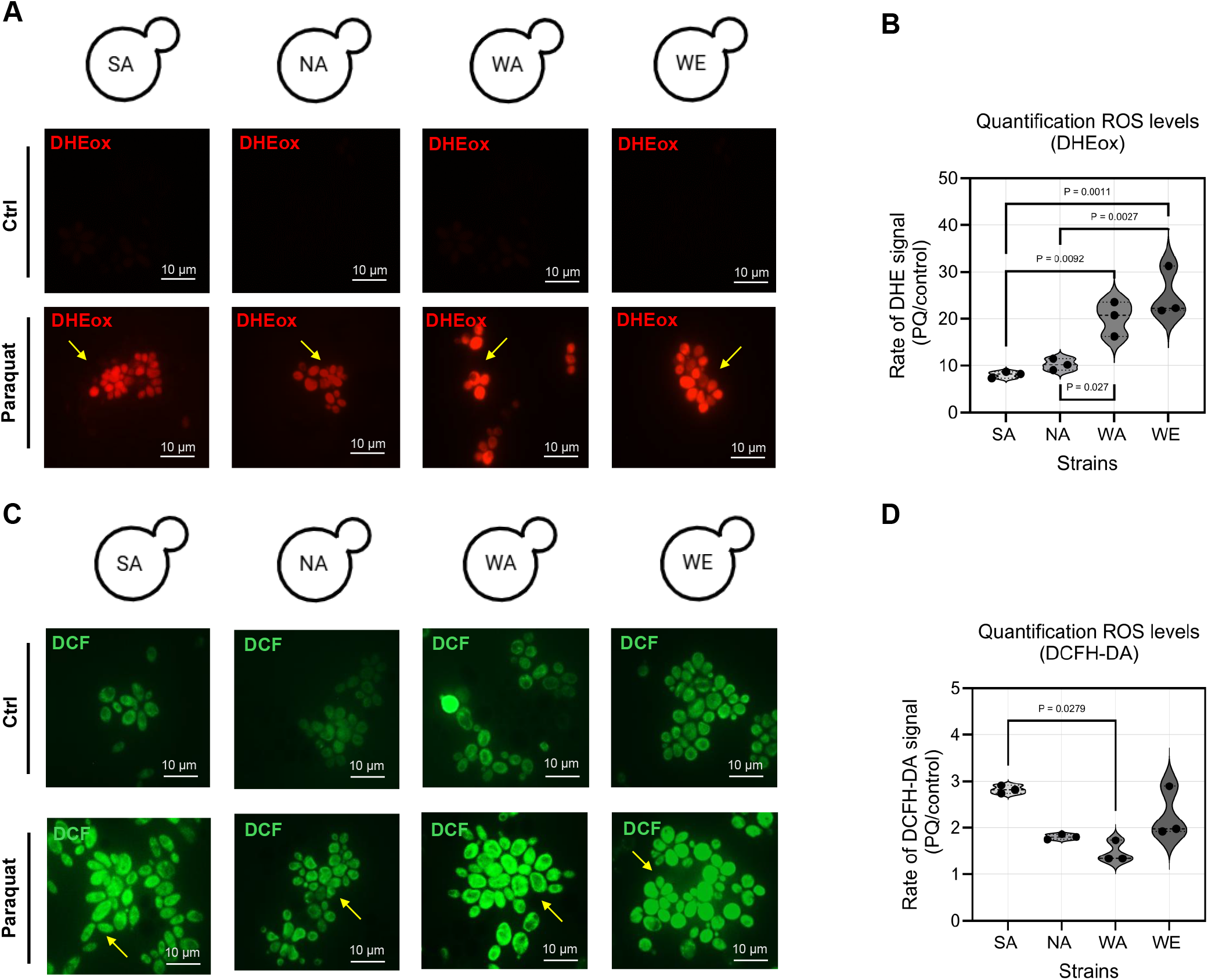
Cellular response of *S. cerevisiae* to oxidative stress (superoxide anion) induced by PQ exposure. (A) Confocal images of DHE staining in untreated PQ-exposed (75 μg/mL) SA, NA, WA, and WE yeast strains at 63x objective and 5x zoom. Yellow arrows: red fluorescence marks the probe oxidized by reactive molecules (DHEox). (B) Fold change (PQ/control) of ROS (DHE) levels in all strains obtained from the quantification of confocal micrographs. (C) Confocal images of DCFD-DA staining in untreated and PQ-exposed of SA, NA, WA, and WE yeast strains visualized at 63x objective and 5x zoom. Yellow arrows: green fluorescence in all strains when exposed to PQ. (D) Fold change (PQ/control) of ROS (DCF) levels from the quantification of confocal micrographs. Biological replicates “N=3”. Values are mean ± SE, Shapiro Wilk normality test p-value > 0.05, ANOVA test p-value < 0.05, post-hoc Tukey. p-value ≤ 0.05 (*), p-value < 0.01 (**), p-value < 0.001 (***), p-value < 0.0001 (****), ns = not significant (p-value > 0.05). Statistical analysis was performed in SPSS software version 20.0 and plotted in GraphPad and RStudio.

### PQ induced strain-specific vacuolar adaptations

To investigate how yeast strains adapt to PQ exposure, we examined their vacuolar morphology, as changes in vacuole structure often indicate activation of stress response mechanisms (Saldaña et al., 2021). Vacuoles are typically classified into three phenotypes: A, up to three large vacuoles per cell; B, multiple small vacuoles with defined borders, and C, highly fragmented vacuoles (Figure 3A) (Aufschnaiter and Büttner, 2019).

**Figure 3:**
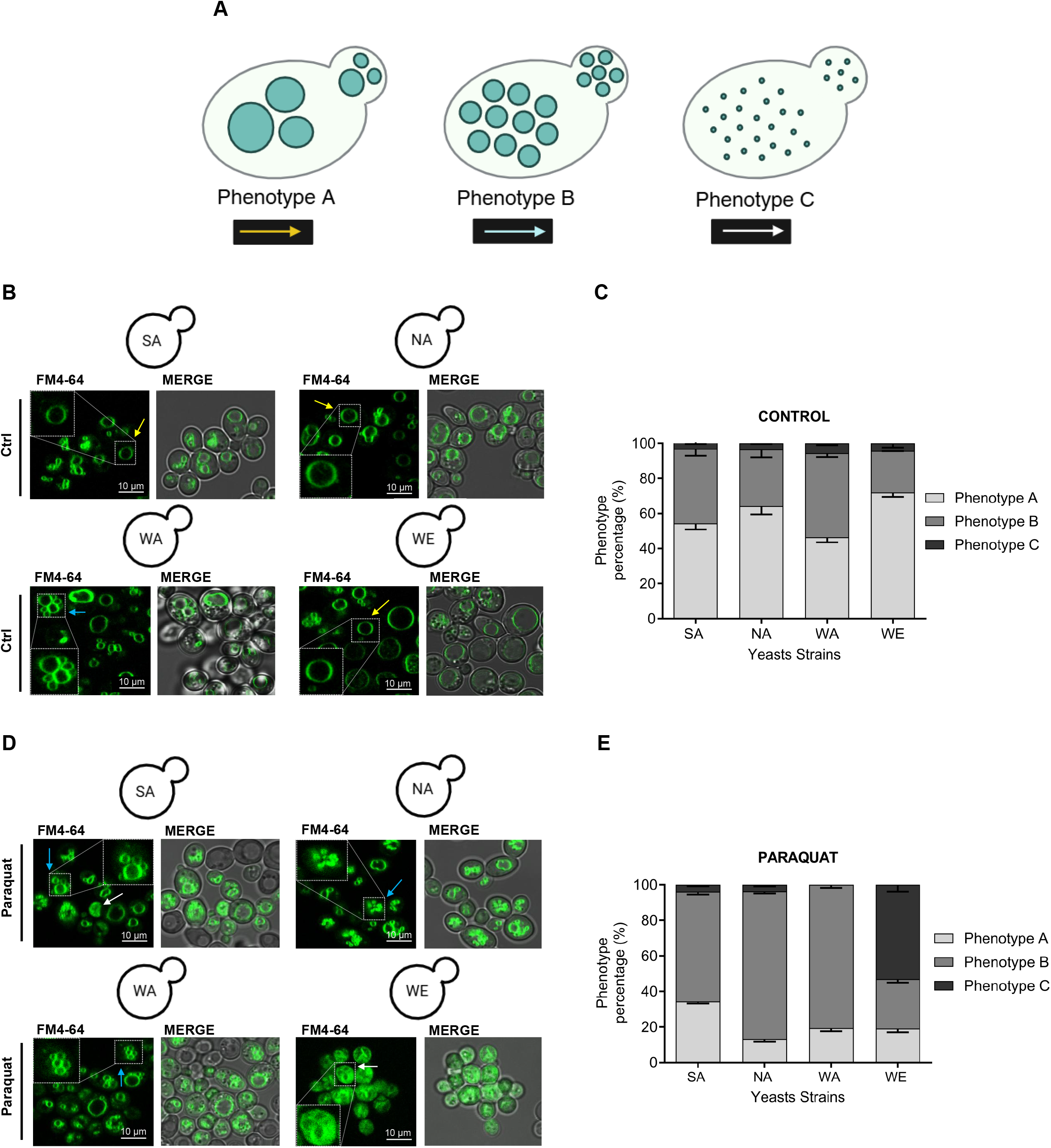
Cellular response of *S. cerevisiae* to PQ-exposure involving vacuolar changes. (A) Diagram of vacuolar forms, categorized by phenotypes A, B, and C. (B) Confocal microscopy of FM4-64 staining of SA, NA, WA, and WE yeast strains in control condition (C) Quantification of vacuolar phenotypes in control condition. (D) Confocal microscopy of FM4-64 staining of SA, NA, WA, and WE yeast strains under PQ treatment. (E) Quantification of vacuolar phenotypes under PQ treatment. Yellow arrows indicate phenotype A; blue arrows phenotype B; and white arrows phenotype C. Biological replicates “N= 3”. Values are mean ± SD, Shapiro Wilk normality test p-value > 0.05, Kruskal Wallis test p-value <0.05. p-value ≤ 0.05 (*), p-value < 0.01 (**), p-value < 0.001 (***), p-value < 0.0001 (****), ns = not significant (p-value > 0.05). Statistical analysis was performed in SPSS software version 20.0 and plotted in GraphPad.

We quantified the percentage of each vacuolar phenotype in four yeast strains under both control conditions and after 48 hours of exposure to 75 μg/mL PQ (Figure 3B-D). Our analysis revealed distinct changes in vacuolar morphology across the strains. Specifically, the prevalence of phenotype A significantly decreased in strains NA (64% control vs. 13% PQ; p = 0.002) and WE (79% control vs. 19% PQ; p = 0.006) following PQ treatment (Figure 3B-E). In contrast, the proportion of phenotype A remained statistically unchanged in strains SA (p = 0.273) and WA (p = 0.184).

Interestingly, strain NA showed a significant increase in phenotype B (a 51% relative increase, from 32% in control to 83% after PQ; p = 0.009) (Figure 3C, 3E). Conversely, phenotype C, representing highly fragmented vacuoles, significantly decreased only in strain WA (a 100% relative decrease, from 6% in control to 0% after PQ; p = 0.016) (Figure 3C, 3E). Microscopic images (Figure 3B, 3D) visually support these quantitative findings, illustrating a reduction in phenotype A and a corresponding rise in phenotype B in the PQ-treated NA strain (Figure 3E) compared to the control (Figure 3C).

The WE strain exhibited a notable increase in phenotype C, rising from 4% in the control condition (Figure 3D) to 53% after PQ exposure (Figure 3E) (p = 0.082). Although this change did not reach the conventional threshold for statistical significance, a clear shift in vacuolar morphology towards fragmentation is evident in the micrographs (Figure 3B and 3D). Compared to the control, the PQ-treated WE strain displayed a marked decrease in phenotype A and a substantial increase in phenotype C. This transition towards a fragmented vacuolar state (phenotype C) in the WE strain upon PQ exposure may indicate a stress response mechanism aimed at preserving organelle integrity to enhance survival.

### Superoxide levels correlate with physiological growth across PQ-treated yeast strains

To elucidate the compensatory mechanism to PQ exposure, we examined the relationships between the different measured traits through correlations. Multiple correlations were performed between PQ-induced fold change in μMax, DHE and DCFH-DA probes, and vacuolar morphologies A, B and C. Most traits were not interconnected (Pearson correlations) (Figure 4A-B). However, a significant negative correlation (Pearson correlation: r = -0.97, p-value = 0.032) emerged between changes in superoxide anion levels and reproductive fitness (Figure 4A-B), suggesting a shared adaptive response to PQ exposure involving intracellular ROS levels and the ability of yeasts to reproduce.

**Figure 4:**
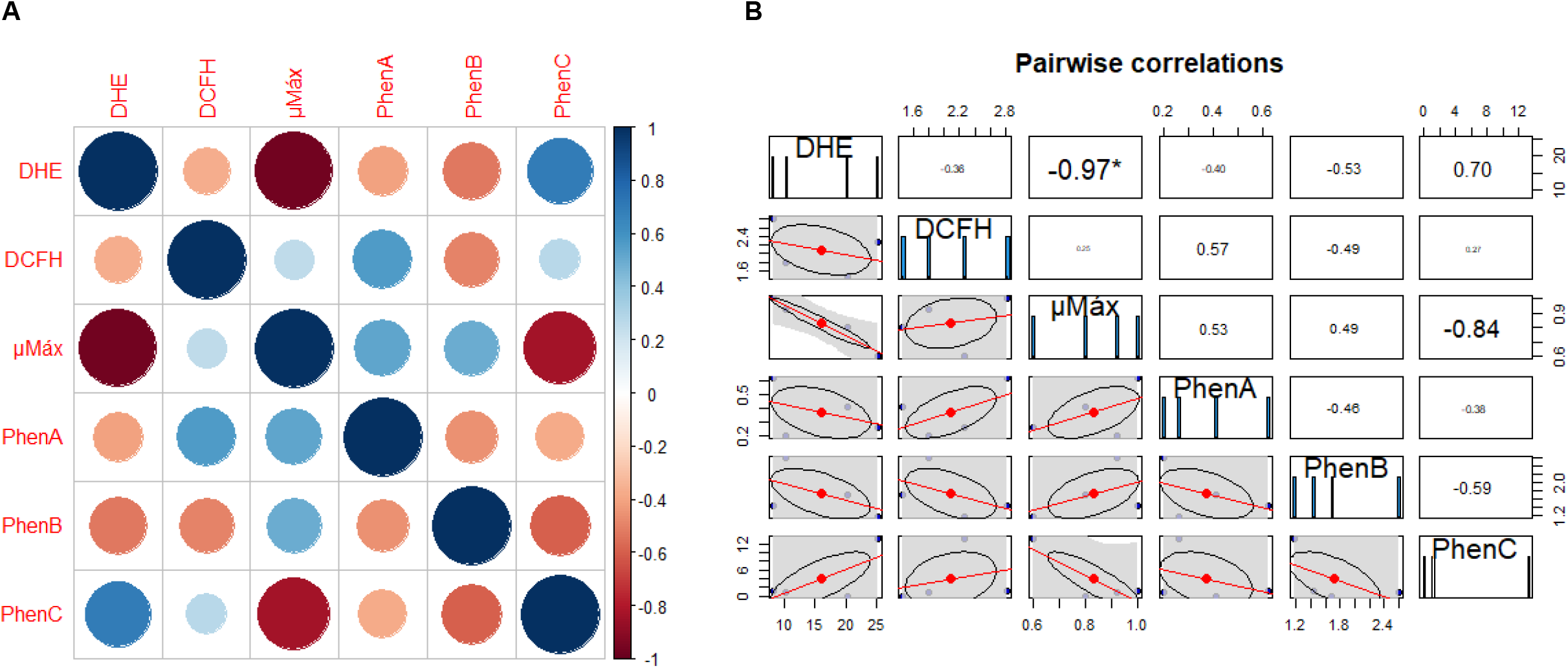
Interplay between physiological parameters, oxidative stress and vacuolar morphology. (A) Plot of multiple correlations between fold change of μMax, DHE probe, DCFH-DA probe, phenotype A, phenotype B and phenotype C. (B) Pairwise correlations between fold change of μMax, DHE probe, DCFH-DA probe, phenotype A, phenotype B and phenotype C. Biological replicates “N=4”. Values are mean ± SE, Pearson correlation p-value < 0.05 in μMax, DHE probe, DCFH-DA probe, phenotype A and phenotype B. For phenotype C and combinations with other parameters were used Spearman correlation. Shapiro Wilk test p-value > 0.05. p-value ≤ 0.05 (*), p-value < 0.01 (**), p-value < 0.001 (***), p-value < 0.0001 (****), ns = not significant (p-value > 0.05). Statistical analysis was performed in SPSS software version 20.0 and plotted in RStudio.

### Genetic determinants of phenotypic responses to PQ exposure

We selected SA, NA, WA, and WE strains for this study because their genomes are fully sequenced (Liti et al., 2009), facilitating the association of gene variants with phenotypes. Considering the established roles of genes involved in lysosomal function and PD development, including humans exposed to pesticides (Ngo et al., 2024), we investigated genomic variations in this pathway across the four yeast strains. We analyzed 41 lysosomal/vacuolar genes, of which only 28 had orthologs in yeast. In these 28 vacuolar genes, we identified a total of 954 predicted variants, comprising 23.7% nonsynonymous, 76.1% synonymous, 0.1% frameshift deletion, and 0.1% start-loss variants (Supplementary Table 1). We hypothesized that missense variants, which are likely to impact protein function, could contribute to PQ responses. To explore this, we performed Pearson’s correlations between the number of deleterious variants as predicted by SIFT (Sorting Intolerant From Tolerant) in vacuolar genes with multiple phenotypes: μMax, DHE (superoxide levels), and vacuolar phenotypes across four yeast strains. Only the statistically significant associations were listed in Table 1.

**Table 1:** Correlations of lysosomal and oxidative stress genes with the number of variants in yeast strains. The potential effects of variants in each gene of the four yeast strains were determined using SIFT. The number of variants in oxidative stress and lysosomal genes was correlated with μMáx, ROS levels (DHE probe), and phenotype B using Pearson correlations. To adjust for multiple comparisons, we applied the Bonferroni procedure to control for the error rate and establish a corrected statistical significance threshold and adjusted P-value (*P*_*adj*_ *a*) based on a significance level of 0.05. Statistical analysis was performed in RStudio.

|  |  | Number of variants |  |  |  | μMax |  |  | DHE |  |  | Phenotype B |  |  |
| --- | --- | --- | --- | --- | --- | --- | --- | --- | --- | --- | --- | --- | --- | --- |
| Human lysosomal genes | Orthologue in *S. cerevisiae* | SA | NA | WA | WE | *P-value* | *P<sub>adj a</sub>* | *r* | *P-value* | *P<sub>adj a</sub>* | *r* | *P-value* | *P<sub>adj a</sub>* | *r* |
| *CLN3* | *YHC3* | 5 | 4 | 3 | 9 | 0.438 | 1.000 | -0.562 | 0.489 | 1.000 | 0.511 | **0.025** | 0.300 | **-0.975** |
| *LIPA* | *YEH1* | 0 | 3 | 6 | 11 | **0.020** | 0.240 | **-0.980** | 0.745 | 1.000 | -0.256 | 0.368 | 1.000 | -0.633 |
| *SLC17A5* | *FEN2* | 6 | 7 | 8 | 1 | 0.398 | 1.000 | 0.601 | 0.528 | 1.000 | -0.472 | **0.026** | 0.312 | **0.974** |
| *ABHD5* | *ICT1* | 8 | 10 | 12 | 3 | 0.540 | 1.000 | 0.460 | 0.400 | 1.000 | -0.600 | **0.032** | 0.384 | **0.968** |
| *ATG5* | *ATG5* | 6 | 7 | 6 | 3 | 0.197 | 1.000 | 0.803 | 0.659 | 1.000 | -0.341 | **0.048** | 0.576 | **0.952** |
| *HSPA8* | *SSA2* | 32 | 27 | 27 | 31 | 0.965 | 1.000 | 0.035 | **0.028** | 0.336 | **0.972** | 0.255 | 1.000 | -0.745 |
|  |  | 87 | 96 | 91 | 65 | 0.273 | 1.000 | 0.727 | 0.565 | 1.000 | -0.436 | **0.018** | 0.216 | **0.982** |
| PQ: μMáx (OD/h) |  | **0.26** | **0.24** | **0.18** | **0.14** |  |  |  |  |  |  |  |  |  |
| PQ: DHE (A.F.U) |  | **23668** | **15665** | **17878** | **25999** |  |  |  |  |  |  |  |  |  |
| PQ: Phenotype B (%) |  | **61.7** | **83.0** | **80.6** | **27.8** |  |  |  |  |  |  |  |  |  |

The number of *YEH1* variants showed a significant negative correlation with μMax (Pearson’s r = - 0.980, p-value = 0.020). A significant positive correlation was found between *SSA2* variant count and DHE-measured ROS levels (r = 0.972, p-value = 0.028). Phenotype B was positively correlated with variants in *FEN2*, while an unexpected negative correlation was observed with variants in *YHC3* (Table 1). However, after adjusting for multiple comparisons (see “Experimental procedures”), these correlations did not remain significant when considering all nonsynonymous and synonymous variants within the gene set. This loss of significance was likely due to the conservative nature of the Bonferroni correction. Consequently, we decided to functionally validate one of the identified genes.

### *fen2*-deletion increases vacuolar fragmentation upon PQ exposure

*FEN2* encodes a proton-pantothenate symporter, and its human ortholog, *SLC17A5*, is a lysosomal protein associated with sialic acid storage disorder and considered a potential susceptibility factor for PD. Given the strong correlation observed between the number of *FEN2* variants and the percentage of vacuolar B phenotype, we investigated the effect of *FEN2* deletion on PQ-induced vacuolar disaggregation. Vacuolar morphology was classified into phenotypes A, B, and C in both the wild-type (BY4742 strain) and the *Δfen2* strain under control (Figure 5A, 5B) and following PQ exposure (200 μg/mL for 48 hours) (Figure 5C, 5D). The percentage of each phenotype was quantified (Figure 5B, 5D). Under control conditions, the *Δfen2*, showed a significant increase in the percentage of phenotype B (p-value = 0.0313, T-test) and C (p-value = 0.0204, T-test), along with a decrease in the percentage of phenotype A (p-value = 0.0114, T-test) compared to BY4742 (Figure 5B). Notably, the decrease in phenotype A of Δfen2 was amplified upon PQ exposure (p-value = 0.0478, T-test). In addition, we found an increased effect of PQ on phenotype C of Δfen2 (p-value = 0.0040, T-test). Specifically, PQ-treated *Δfen2* strain exhibited a more pronounced increase in the percentage of phenotype B (p = 0.0270, t-test) and C (p = 0.0071, t-test), and a more pronounced decrease in the percentage of phenotype A (p = 0.0051, t-test) compared to BY4742 (Figure 5D). These findings suggest a potential role for FEN2 in mediating the vacuolar response to PQ.

**Figure 5:**
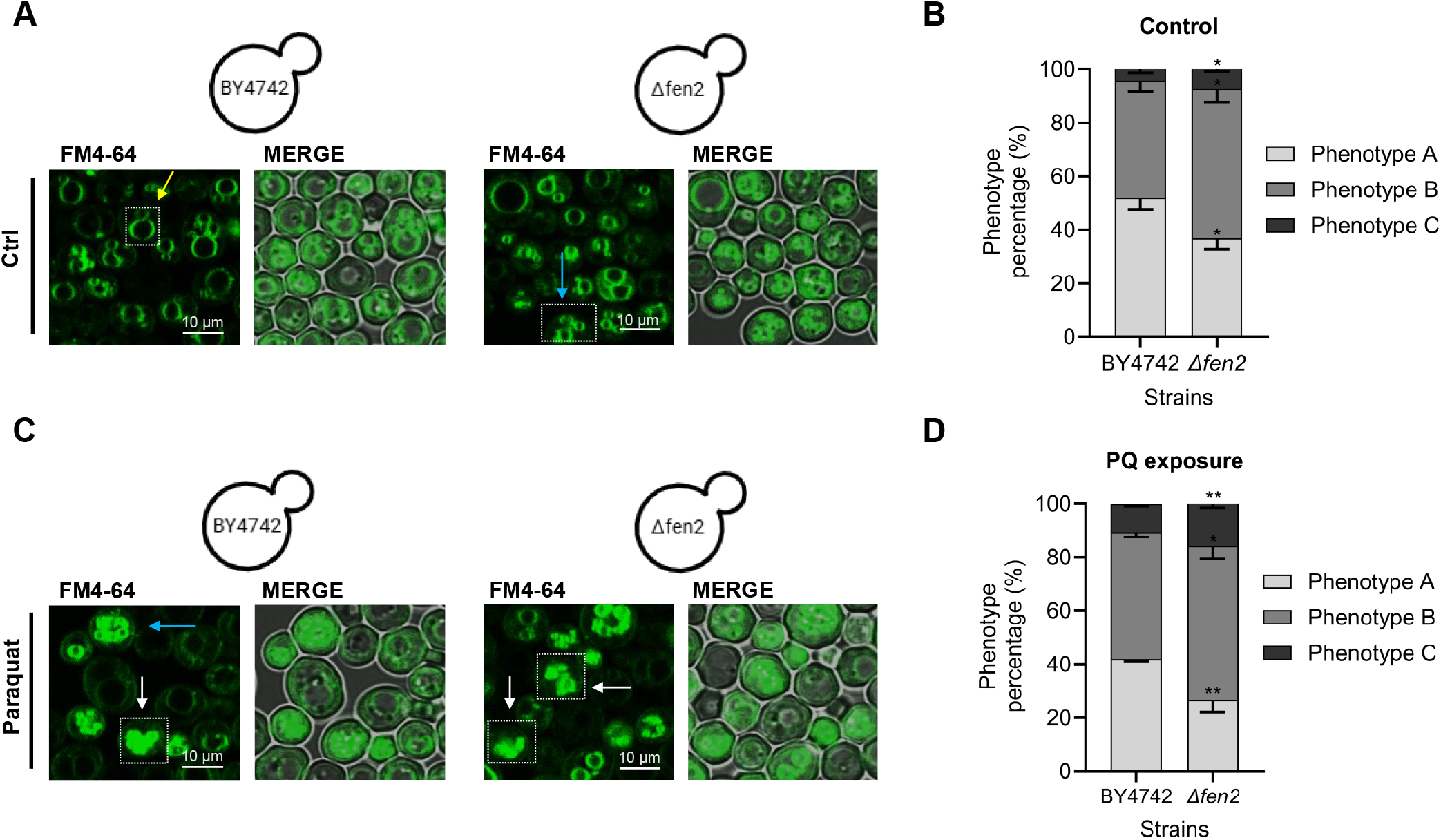
Cellular response of BY4742 (ref) and *fen2*-deletant strain to PQ exposure involving vacuolar changes: (A) Confocal microscopy of FM4-64 staining in BY4742 and *fen2*-deletant (*Δfen2*) control condition. (B) Quantification of vacuolar phenotypes in control condition. (C) Confocal microscopy of FM4-64 staining in BY4742 and *fen2*-deletant (*Δfen2*) under PQ treatment (200 μg/mL). (D) Quantification of vacuolar phenotypes under PQ treatment. Yellow arrows: phenotype A, blue arrows: phenotype B, and white arrows: phenotype C. Biological replicates “N= 3”. Values are mean ± SD, Shapiro-Wilk normality test p-value > 0.05, T-test p-value <0.05. p-value ≤ 0.05 (*), p-value < 0.01 (**), p-value < 0.001 (***), p-value < 0.0001 (****), ns = not significant (p-value > 0.05). Statistical analysis was performed in SPSS software version 20.0 and plotted in GraphPad.

## Discussion

In humans, PQ is highly toxic (Sukumar et al., 2019) with potential absorption through the gastrointestinal tract, skin contact, or inhalation (Shi et al., 2023). Exposure can induce several pathologies, including lung fibrosis, liver tumors, and PD (Donaher and Van den Hurk, 2023). PQ disrupts the intracellular redox cycle (Donaher and Van den Hurk, 2023), leading to mitochondrial fragmentation via the generation of ROS and reactive nitrogen species, as well as lysosomal damage (Pascua-Maestro et al., 2017). These cellular injuries can trigger apoptosis in dopaminergic neurons (Alural et al., 2015; Xiong et al., 2019), contributing to PD pathogenesis (See et al., 2022). Given that these pathways are conserved in eukaryotes, *S. cerevisiae* has been a valuable model for elucidating PQ toxicity mechanisms (Sillapawattana et al., 2024). However, the majority of these investigations have focused on single yeast strains, limiting our understanding of how genetic diversity influences the cellular and molecular adaptive strategies to PQ.

Our study demonstrates a differential impact of PQ on μMax across four genetically diverse *S. cerevisiae* strains. Under physiological conditions, μMax is controlled by GCN4, a transcriptional factor that regulates amino acid biosynthesis, along with other factors as FIL1 and GCN2 (Duncan et al., 2018; Fendt et al., 2010). Whether changes in amino acid homeostasis relate to the strain-specific variations in μMax upon PQ exposure has to be determined. Our results highlight the significance of genetic background in determining yeast susceptibility to PQ toxicity, which is consistent with previous findings in other models (Lovejoy et al., 2021; Yin et al., 2011).

Oxidative damage is a PD hallmark. Notably, there is a correlation between the levels of oxidative stress markers and PD age of onset and severity in both blood and cerebrospinal fluid (Ihara et al., 1999; Takahashi et al., 2021). Consistent with these observations, our findings indicate that strains exhibiting high superoxide levels show reduced reproductive fitness (a susceptibility phenotype), implying less efficient mechanisms for mitigating oxidative stress. This finding highlights the value of yeast of different backgrounds for precision medicine studies and suggests that strategies aimed at reducing oxidative damage could be useful to prevent PQ-induced cell damage in humans.

Superoxide is a common product of electron leakage from donor redox centers within the mitochondrial electron transport chain. This radical species can lead to the formation of secondary products such as peroxide, a reaction catalyzed by mitochondrial superoxide dismutase. Unexpectedly, our results revealed no significant correlations between peroxide levels and any of the parameters measured, suggesting that superoxide dismutase activity varies across the yeast strains. In the PQ-treated cells, superoxide may directly damage lipids, proteins, and DNA (Kowaltowski and Vercesi, 1999) or exert its effects in combination with other radical species, including peroxynitrite.

In yeast, vacuoles, the functional counterparts of mammalian lysosomes, are highly dynamic organelles that exhibit a continuous cycle of fusion and fission in response to environmental cues (Aufschnaiter and Büttner, 2019; Li and Kane, 2009). Notably, oxidative stress is a recognized inducer of vacuolar fragmentation (Corson et al., 1999; Kim et al., 2021). This process is under the stringent control of vacuolar protein sorting 1 (VPS1) and autophagy-related protein 8 (ATG8) (Mikawa et al., 2010). VPS1, a protein implicated in intracellular trafficking, is also essential for the development of cellular resistance to oxidative insults. Concurrently, ATG8 contributes to the structural integrity of the vacuolar membrane, independent of its canonical role in autophagy (Ishii et al., 2019; Mikawa et al., 2010). Double mutants in *Vps1* and *Atg8* exhibit enhanced sensitivity to PQ, osmotic stress, and calcium overload than the single mutations, indicating impaired vacuolar function (Mikawa et al., 2010). Further investigation is required to elucidate the specific role of these proteins in determining the spectrum of PQ susceptibility observed across diverse yeast strains.

Genetic studies demonstrate a direct association between various mitochondrial and lysosomal genes and the risk of idiopathic PD (iPD) (Blauwendraat et al., 2020). Among the highest PD risk factors is the *GBA1* gene, which encodes for lysosomal β-glucocerebrosidase (GCase) (Sidransky et al., 2009). Remarkably, GCase plays roles in both organelles: in the lysosome, degrading glucosylceramide, and mitochondria, promoting complex I stability (Klein and Outeiro, 2023; Rubilar et al., 2024). Since pesticides target mitochondria, a recent study identified enrichment in genomic variants in 26 genes associated with lysosomal function. These results support the idea that impaired lysosomal and mitochondrial dysfunction is necessary to develop PD (Ngo et al., 2024). Similarly, our yeast study shows correlations between the number of missense variants in different genes associated with vacuolar/lysosomal function and PQ-induced phenotypes. Our analysis revealed a positive association between an increased number of variants in the *FEN2* gene with a rise in phenotype B. In yeast, the *FEN2* gene coregulates carbon and nitrogen catabolism, and amino acids and ergosterol biosynthesis (Marcireau et al., 1996; Stolz and Sauer, 1999). The human orthologue of this gene belongs to the SLC17A transport family (Cao et al., 2014), and is implicated in the sialuria, a sialic acid lysosomal storage disorder with enlarged lysosomes (Huizing et al., 2021; Tarailo-Graovac et al., 2017). While in 2017 an exome sequencing study identified *SLC17A5* as a potential PD susceptibility gene (Robak et al., 2017), a recent 2025 larger study focused on European ancestry, found no association between *SLC17A5* common variants and PD risk, potentially due to limitations to detect rare variants, and the exclusion of non-European populations. This discrepancy highlights the need for comprehensive genetic analyses across diverse populations, like our study in yeast in of different genetic backgrounds, to fully understand the role of sialic acid homeostasis in different PD subtypes associated with lysosomal dysfunction (Sabir et al., 2025). Our results showed significantly increased vacuolar fragmentation in a *fen2*-deletant strain, likely due to the accumulation of a sialic acid-like product in yeast. This may reflect the sialuria phenotype in humans, where lysosomal dysfunction from transport deficiency leads to similar fragmentation and could be implicated in pesticide-induced PD.

Our study also has limitations: i) the use of a small number of strains; which limits the power to identify significant associations across traits; ii) the focus on only on two PD-related phenotypes: ROS and vacuolar adaptations, omitting others like proteostasis (Kulkarni et al., 2023); iii) the use of domesticated yeast strains which while offering genetic stability, may not fully represent the genetic and phenotypic variability found in wild populations: and iv) the phenotypic characterization, lacking mechanistic assays; among others. However, by investigating the cellular and molecular responses underlying resistance and susceptibility to PQ across these diverse genetic backgrounds, our study reveals the biological foundation supporting these differences and highlights the fundamental role of genetics in understanding these complex processes.

In conclusion, our work explores cellular and molecular mechanisms underlying PQ toxicity in four different yeast backgrounds, which resulted in good proxies for studying PQ susceptibility/resistance in humans. Furthermore, we validated the association between superoxide levels and reproductive fitness and the role of *FEN2* in PQ-induced vacuolar disaggregation. Understanding the biology of the PQ-resistant strains may facilitate the design of strategies to prevent pesticide-induced PD. These can be achieved by gene mapping studies and omics in *S. cerevisiae*, followed by validations in animal models and human cells.

## Experimental procedures

### Yeast strains and culture conditions

The four haploid (*MAT α* and *MAT a*) *S. cerevisiae* strains used in this study were: North American (NA; YPS128, *MAT α/a, ho: HygMX, ura3::kanMX*), West African (WA; DBVPG6044, *MAT α/a, ho: HygMX, ura3::kanMX*), Sake (SA; Y12, *MAT α/a, ho: HygMX, ura3::kanMX*) and. Wine/European (WE; DBVPG6765, *MAT α/a, ho: HygMX, ura3::kanMX*) (Brice et al., 2018; Salinas et al., 2016). These strains were stored in solid media (YPD) at 4°C. Liquid and solid culture media were used depending on the experiments to be performed: YPD media (2% glucose, 2% peptone, 1% yeast extract) and YNB (0.67% YNB base without amino acids, 0.2% uracil, 0.0875% com drop out and 2% glucose). Validation studies were performed using the BY4742 strain, specifically the wild-type (WT; Accession Y10000), the *fen2*-deletant (BY4742; MATα; his3Δ1; leu2Δ0; lys2Δ0; ura3Δ0; YCR028c::kanMX4). Reference and deletant strain were obtained from Euroscarf (http://www.euroscarf.de).

### Growth curves conditions

Yeast cells were pre-cultured in 200 μL of YNB medium supplemented with uracil (0.2% uracil) for 48 h at 28 °C. For the experimental run, the four yeast strains were inoculated to an optical density (OD) of 0.03-0.1 (wavelength of 620 nm) in 200 μL of medium and incubated without shaking at 28 °C for 48 h (YNB control and Paraquat (Sigma Aldrich-Merck, CAS No. 75365-73-0) at 75 μg/mL) in a Tecan Sunrise absorbance microplate reader (Tecan Trading AG, Männedorf, Switzerland). OD was measured every 30 min using a 620 nm filter. Each experiment was performed in triplicate. Growth rates for each strain were calculated as previously described (García-Ríos et al., 2014; Quispe et al., 2017). Briefly, OD measurements as a function of time were fitted to the mathematical model of the re-parameterized Gompertz sigmoid curve describing microbiological temporal growth, previously proposed (Zwietering et al., 1990) from which growth rates were obtained. The parameters of each curve were processed in the GrowthRates software to obtain OD max, μmax, and the average time of the lag phase. Finally, the area under the curve (AUC) values of the growth curves were calculated using R commands.

### Measurement of levels of Reactive Oxygen Species (ROS)

The yeast strain cultures were incubated for 48 hours at 28°C in 3 ml of YNB medium and centrifuged at 1700 x g at 4 °C for 4 min. Intracellular levels of superoxide anion were detected through the dihydroethidium (DHE) probe (Thermofisher Scientific, NºD1168, U.S.A), and intracellular levels of hydrogen peroxide (H_2_O_2_) were detected through the 2′,7′-dichlorofluorescein diacetate (DCFH-DA) probe (Sigma Aldrich, NºD6883, U.S.A). The pellet was incubated with the DHE probe at a concentration of 10 μg/mL (Mendes-Ferreira et al., 2010) in 500 μL of PBS (80 mM Na_2_HPO_4_, 20 mM NaH_2_PO_4_ and 100 mM NaCl) (Rego et al., 2020) for 30 min and washed with PBS as previously reported (Mendes-Ferreira et al., 2010; Rego et al., 2020). The DCFH-DA probe was used at a concentration of 10 μM (Chapela et al., 2022) in 500 μL of PBS for 1 h and washed with a PBS buffer. Fluorescence intensity was visualized using a Leica SP8 confocal microscope (Leica Microsystems, Germany) at 518/605 nm for DHE and 490/530 for DCFH-DA. For microscopic analysis, 4 μL of each incubated strain (DHE and DCFH-DA probe) were taken, added to a slide, and squeezed with a coverslip gently for *in vivo* visualization and observed under a microscope using a 63X objective and 5X magnification. Images were processed in ImageJ (Schneider et al., 2012) and LASX software (Leica Microsystems, Germany).

### FM4-64 internalization assay in *S. cerevisiae*: Vacuolar phenotypes

Yeasts were seeded on a plate with YPD medium and 2% agar-agar. Isolations were made from each strain to obtain single colonies previously incubated at 28°C without shaking for 48 hours. Then, inocula were transferred into 500 μL of liquid YPD and incubated with 3 μM of FM4-64 (Ex: 565 nm - Em: 744 nm) (ThermoFisher Scientific, N° F34653, U.S.A) for 24 hours in 48-hour cultures in control condition (Seeley et al., 2002) and with 75 μg/mL Paraquat at 28°C with shaking (150 rpm) in the dark. 4 μL of each incubated strain was added to a slide and squeezed with a coverslip gently for in vivo viewing and observed on Leica SP8 confocal microscope (Leica Microsystems, Germany), using 63X objective and 5X magnification. Images were processed in ImageJ and LASX software (Leica Microsystems, Germany). The vacuolar characteristics of 110-150 cells per field were analyzed and classified according to their phenotype into “A”, “B” or “C”, as proposed by Seeley E. et al. (2002) (Seeley et al., 2002). The assay was repeated three times and quantified by three researchers independently for each yeast strain.

### Statistical analysis

Growth parameters μMax, ROS level by DHE and DCFH-DA, and percentage of vacuolar phenotypes were used as quantitative variables in the control condition and PQ at a 75 μg/mL concentration. For reference and deletant strain, we used a PQ concentration at 200 μg/mL. Shapiro-Wilk normality test p>0.05, parametric ANOVA test p<0.05 with multiple comparisons, and non-parametric Kruskal-Wallis test p-value < 0.05 with multiple comparisons were used for all measurements. For the percentage of vacuolar phenotypes in deletant strains validation, the Shapiro-Wilk normality test (p>0.05) and T-test (p<0.05) were done. Sample “N” in the experiments was 5 biological replicates and 3 technical replicates per sample for μMax, 6 biological replicates and 3 technical replicates per sample for ROS level by DHE and DCFH-DA probe, and 3 biological replicates with 3 technical replicates per sample for quantification of vacuolar phenotypes. Statistical significance was considered for a value of p<0.05. Furthermore, multiple correlations were conducted in this study. Pearson correlations were performed for the fold change of DHE, DCFH, μMax, and vacuolar phenotypes A, B and C, as the data exhibited normal distribution (Shapiro-Wilk test p > 0.05). The potential effects of variants in each gene of the four yeast strains were determined using SIFT software (Table 2) and the *S. cerevisiae* S288C reference genome (https://www.yeastgenome.org/). The number of variants in oxidative stress and lysosomal genes was correlated with μMáx, ROS levels (DHE probe), and phenotype B using Pearson correlations. To adjust for multiple comparisons, we applied the Bonferroni procedure to control for the error rate and establish a corrected statistical significance threshold and adjusted P-value (*P*_*adj*_ *a*) based on a significance level of 0.05. Statistical analysis was performed in SPSS software version 20.0 (Illinois, U.S.A) and plotted in GraphPad (California, U.S.A) and RStudio (U.S.A).

## Acknowledgements

This work was supported by ANID-CHILE: Fondecyt grant No 1230317 (A.D.K)

## CRediT author statement

JCR: Conceptualization, Methodology, Investigation, Data Analysis, Data Curation, Writing-Original draft preparation, and Editing. BZ: Methodology, Data Analysis. FAC: Conceptualization, Methodology, Supervision, Writing-Reviewing, and Editing. ADK: Conceptualization, Methodology, Data curation, Writing-Original draft preparation, Supervision, Writing-Reviewing, Editing, and Funding the Project.

## Figure legends

**Supplementary Table 1: SIFT-predicted variants in vacuolar genes in the four yeast strains**. The first sheet shows the human lysosomal genes and their yeast orthologues. The second sheet shows the reference allele (Ref allele), and reference amino acid (Ref amino acid), type of predicted variants, and SIFT predictions across SA, NA, WA, and WE yeast strains.

